# Early frameshift alleles of zebrafish *tbx5a* that fail to develop the heartstrings phenotype

**DOI:** 10.1101/103168

**Authors:** Elena Chiavacci, Lucia Kirchgeorg, Anastasia Felker, Alexa Burger, Christian Mosimann

**Affiliations:** Institute of Molecular Life Sciences, University of Zürich, Winterthurerstrasse 190, 8057 Zürich, Switzerland

**Keywords:** tbx5a, Tbx5, zebrafish, mutagenesis, mutants, CRISPR-Cas9, genetics, heart, forelimbs, Holt-Oram Syndrome

## Abstract

Tbx5 is a key transcription factor for vertebrate heart and forelimb development that causes Holt-Oram syndrome when mutated in humans. The classic zebrafish mutant for *tbx5a* named *heartstrings (hst)* features recessive absence of pectoral fins and a spectrum of heart defects, most-prominently featuring the name-giving stretched heart tube. The mutation of the *hst* allele is a stop codon that is predicted to result in a truncated Tbx5a protein that might feature residual activity. Here, using CRISPR-Cas9 mutagenesis, we generated zebrafish strains for two new *tbx5a* frameshift alleles in the first coding exon: *tbx5a c.21_25del* and *tbx5a c.22_31del,* abbreviated as *tbx5a*^*Δ5*^ and *tbx5a*^*Δ10*^. Homozygous and trans-heterozygous combinations of these new *tbx5a* alleles cause fully penetrant loss of pectoral fins and heart defects including changes in cardiac marker expression akin to *hst* mutants. Nonetheless, distinct from *hst* mutants, homozygous and trans-heterozygous combinations of these *tbx5a* frameshift mutants do not readily manifest the stretched *hst* heart phenotype. Our observation points out the importance and value of comparing phenotypes from different classes of mutant alleles per gene.

## INTRODUCTION

The T-box transcription factor Tbx5 is expressed in the anterior lateral plate mesoderm (ALPM) and contributes to cardiac and forelimb formation^1–3^. Mutations in the human *TBX5* gene cause the autosomal-dominant Holt-Oram Syndrome (HOS) with concomitant heart and arm malformations^4,5^ that *Tb*x5-mutant mice recapitulate^2,3^.

Zebrafish homozygous for the *tbx5a* allele *heartstrings (hst^m^*^21^, or short *hst)* and morpholino-mediated *tbx5a* knockdown mimic HOS phenotypes with defects in heart and pectoral fin formation^6, 7^. Most-prominently, *hst* embryos form a stretched heart tube that inspired the mutant’s name. Nonetheless, while the molecular heart and fin phenotypes are robust, the heartstrings phenotype is variable with penetrance and expressivity linked to the genetic background^7^.

The molecular lesion in *hst* is an ENU-induced stop codon in the second-last coding exon; theoretically, *hst* mRNA can translate into a C-terminally truncated Tbx5a protein with residual or dominant-negative activity^5,7–9^. Similarly, *tbx5a* morpholino knockdown causes the heartstrings phenotype with variable penetrance^6,7,10^. Here, using CRISPR-Cas9 we generated new mutant *tbx5a* alleles with frameshifts in the first coding exon. Our alleles cause recessive phenotypes that recapitulate key defects of *hst* mutants, but do not develop the classic heartstrings phenotype. Our observation underlines the importance of allele comparisons in the design and interpretation of genome editing experiments.

## OBJECTIVE

Generation of new frameshift alleles for *tbx5a* in zebrafish with defined molecular lesions. Subsequent phenotypic analysis and comparison to previously reported *tbx*5a-mutant phenotypes in zebrafish.

## RESULTS AND DISCUSSION

To generate putative *tbx5a* null alleles in zebrafish, we employed Cas9 ribonucleoprotein complex (RNP)-mediated mutagenesis using our established sgRNA[*tbx5^ccA^*] that targets the first coding exon (Fig. 1A)^11^. This sgRNA targets the coding sequence in the first coding exon downstream of the conserved translation initiation codon^12^. We targeted the first exon to introduce frameshift and subsequent stop codons early in the open reading frame to avoid potential translation of N-terminal Tbx5a protein remnants that could retain function. Further indicating that targeting this region could result in loss-of-function alleles, the corresponding amino acid sequence is highly conserved between zebrafish and humans (indicating functional conservation) and human HOS patients have been identified with frameshift-introducing nucleotide insertions at similar positions within *TBX5*^13^.

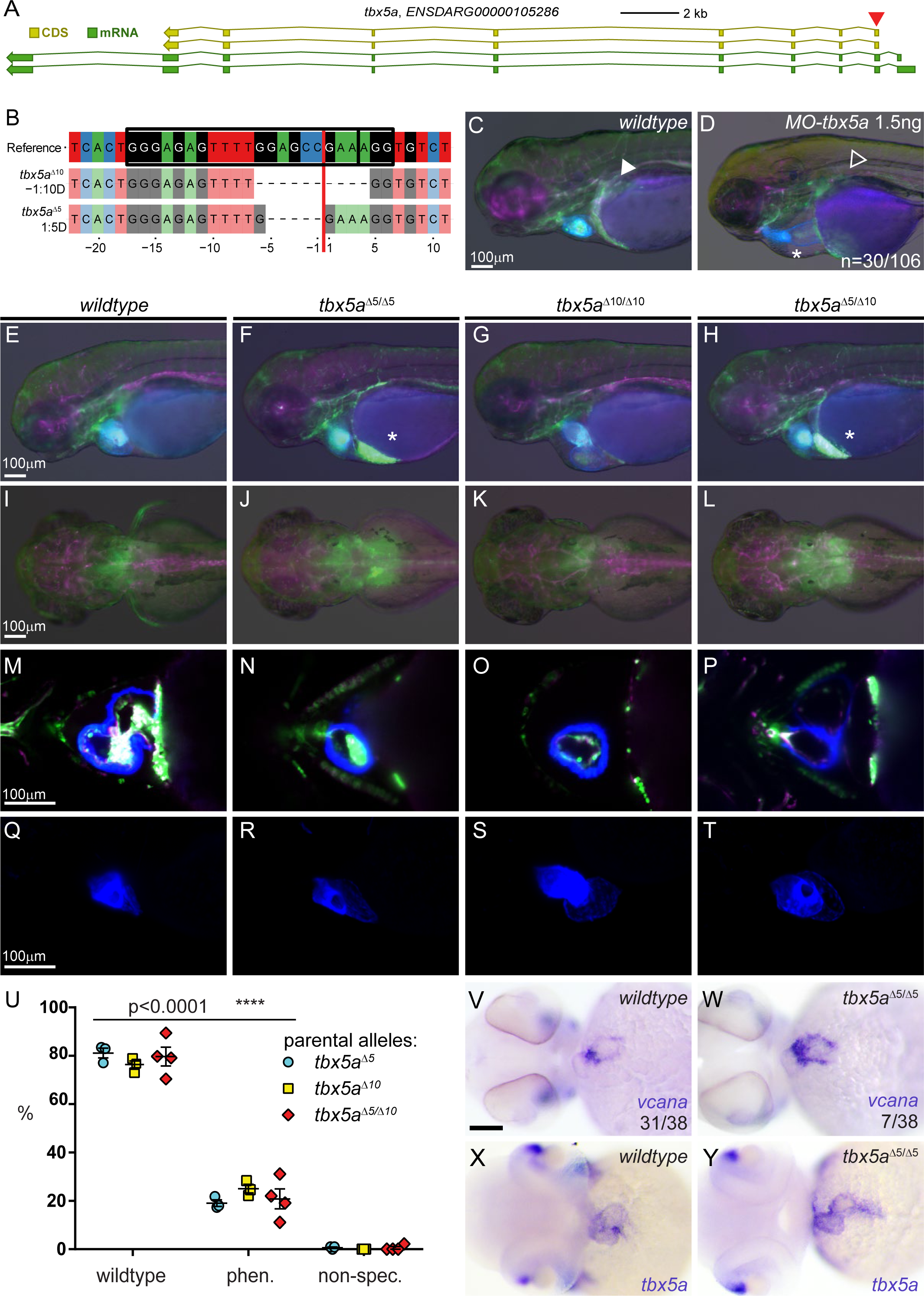
New frameshift alleles in the first coding exon of zebrafish *tbx5a* recapitulate reported mutant phenotypes but fail to develop the heartstrings phenotype. **A**) CRISPR-Cas9-mediated mutagenesis by NHEJ of the first coding exon in *tbx5a.* Gene locus as per genome annotation Zv10 with two major isoforms that share the first coding exon, with red arrowhead showing location of sgRNA used for mutagenesis; orange boxes mark coding exons (CDS), green boxes mark transcribed exons (mRNA). **B**) CrispRVariants panel plot depiction of the isolated germline alleles. Top shows genomic reference, allele *tbx^Δ10^* and *tbx^Δ5^* shown below, resulting in out-of-frame deletions that introduce frameshifts in the coding region. Black boxes over reference sequence indicate sgRNA, smaller box the *5’-NGG-3’* PAM sequence, red line indicates the predicted Cas9-induced double-strand break position. Sequence shown inverse in accordance with figure panel **A**. **C-T**) Comparison of heart and pectoral fin phenotypes from tbx5a morpholino knockdown versus homozygous and trans-heterozygous allele combinations of *tbx5a*frameshift alleles. Images show regions from 72 hpfzebrafish embryos in the tripletransgenic reporter background *RGB (lmo2:dsRED;drl:EGFP,myl7:AmCyan),* anterior to the left, scale bars mark 100μm. **C**,**D**) Translation-blocking morpholino (MO) injection against *tbx5a* causes the heartstrings phenotype with variable penetrance. Compared to wildtype controls (**C**) that form well-looped hearts and pectoral fins (white arrowhead), morpholino-injected embryos (**D)** and miss pectoral fins (open arrowhead) and 28% develop cardiac edema a stretched heart tube (asterisk). **E-H**) Lack of heartstrings phenotypes resulting from recessive *tbx5a*frameshift alleles. Lateral brightfield and fluorescence composite images, white asterisks depict pooling of erythrocytes due to inefficient circulation. Note how all allele combinations (**F-G**) result in hearts with recognizable looping compared to morpholino injected embryos (**D**). **I-L**) Dorsal view, revealing complete lack of pectoral fins in all allele combinations.**M-P**) Ventral view with optical sections taken at the same Z-position from SPIM imaging show in *tbx5a* frameshift mutants the inflated pericardial space surrounding the heart and thinner myocardium, floating ventricles, and looping defects (**N-P**) compared to wildtype (**M**).**Q-T**) Corresponding lateral view of maximum intensity projections from panels **M-P** shown for blue channel *(myl7:AmCyan,* marking cardiomyocytes). **U**) Statistical representation of observed phenotypes shows Mendelian distribution, with no unspecific phenotypes resulting from the mutagenesis or the genetic background. **V,W**) mRNA *in situ* hybridization (ISH) for the cardiac marker *versican a (vcana),* numbers indicate embryos in clutch without prior sorting for phenotypes; *tbx5a^Δ5^* homozygotes show expansion of *vcana* as reported for *hst^m21^* mutants. **X**,**Y**) mRNA*in situ* hybridization (ISH) for *tbx5a* shows the presence of *tbx5a* mRNA in wildtype (**X**) and in homozygous *tbx5a^Δ5^* embryos (**Y**).

Maximized mutagenesis using Cas9 RNPs with sgRNA[*tbx5^ccA^*] cause recognizable *tbx5a* loss-of-function phenotypes in F0 crispants^11^. We injected the sgRNA complexed with Cas9 protein as solubilized RNPs^11^ at sub-optimal concentration to achieve viable mosaicism (see Methods for details) in the multicolor *Tg(lmo2:dsRED2;drl:EGFP;myl7:mCyan)* reporter background, subsequently abbreviated as *RGB.* In *RGB* embryos, dsRED2 labels endothelial, hematopoietic, and endocardial progenitors *(lmo2)* in red^14^, EGFP marks all lateral plate mesoderm lineages *(drl)* including pectoral fins in green^15^, and AmCyan reveals the differentiated cardiomyocytes*(myl7)* in blue^16^; consequently, *RGB* enables *in vivo* imaging of all cardiovascular and additional LPM lineages over the first 3 days of development.

From F0 outcrosses that transmitted mutant *tbx5a* alleles, we genotyped adult F1 zebrafish for the presence of mutated *tbx5a* alleles by tail clipping, PCR, sequencing, and CrispRVariants analysis^17^. From the recovered germline alleles, we kept heterozygous strains for the lesions *c.21_25del* and *c.22_31del* (henceforward abbreviated as *tbx5a^Δ5^* and *tbx5a^Δ10^*) (Fig. 1A, B). These alleles generate out-of-frame mutations starting from base +21 or base +22, respectively, and result in premature stop codons shortly after the conserved initiation codon.

We in-crossed adult F1 heterozygotes for *tbx5a^Δ5^* and *tbx5a^Δ10^* and inter-crossed parents for each allele to assess F2 homozygous and trans-heterozygous embryos for developmental phenotypes at 3dpf. We found that all combinations of the alleles resulted in Mendelian ratios of heart defects (Fig. 1E-H,U) and concomitant, completely penetrant loss of pectoral fins (Fig. 1I-L, U). The cardiac defects for the allele combinations included: cardiac edema with blood accumulation at the inflow tract region (Fig. 1F,H, white asterisks), heart mis-looping (Fig. 1G), and misshapen atrial and ventricular chambers (Fig. 1F-H, N-P, R-T), with n=49/306 for *tbx5a^Δ5^,* n=55/231 for *tbx5a^Δ10^,* n=108/417 for *tbx5a^Δ5/Δ10^* (Fig. 1U). Clutch values including mortality (from now on abbreviated as death rate D.R.) were: for *tbx5a^Δ5^* clutch 1, n=29 D.R.=20.7%; clutch 2, n=144 D.R.=11.1%; clutch 3, n=133, D.R.=24.1%. For *tbx5a^Δ10^* clutch 1, n=78 D.R.=7.7%; clutch 2, n= 65, D.R.=12.3%; clutch 3, n=88, D.R.=0%. For *tbx5a^Δ5/Δ10^* clutch 1, n=58, D.R.=22.4%; clutch 2, n=170, D.R.=3.5%, clutch 3, n=147, D.R.=7.5%.

While cardiac defects were fully penetrant in homozygous and trans-heterozygous mutants, the expressivity of the cardiac phenotype was highly variable, ranging from inflow tract defects (Fig. 1F) to mis-looped chambers (Fig. 1G). Live imaging using selective plane illumination microscopy (SPIM) allowed optical sectioning (Fig. 1M-P) and imaging of the whole heart (Fig. 1Q-T, side view), revealing additional details of the chamber defects. We detected atrium mis-positioning (Fig. 1N,O), freely floating and rounded-up ventricles within the pericardial cavity (Fig. 1N-P, R-T), and thinner cardiac walls (Fig. 1P) compared to wildtype or heterozygous siblings that develop a regularly formed ventricle anchored within the pericardium (Fig. 1M,Q). mRNA expression of *versican a (vcana)* in homozygous *tbx5a^Δ5^* mutants was expanded in tbx5a-mutant hearts (Fig. 1V,W). All these phenotypes are well-documented for both *tbx5a* morphants in which *tbx5a* mRNA is downregulated via morpholino injection^7,9,10,18^ and in embryos homozygous for the classic *tbx5a* allele *hst*^6–8^

Nonetheless, in contrast to the reported morpholino and *hst* mutant phenotypes, we never detected the most-severe form of the *hst* phenotype consisting of a string-shaped heart tube and a deformed head^7^. We readily observed this phenotype using translation-blocking *tbx5a* morpholino injections (n=30/106) (Fig. 1D), in line with previous reports of variable expressivity^7,11,19^. The presence of the *hst* phenotype itself has previously also been linked to the genetic background^7^, suggesting that the *hst* phenotype is a variation of the *tbx5a*loss-of-function phenotype. Taken together, homozygous and trans-heterozygous combinations of our new *tbx5a* frameshift alleles recapitulate morphological and molecular phenotypes of *tbx5a* morphants and the classic *hst* mutant with exception of the heartstrings phenotype. This observation suggests that either our frameshift alleles are not null and possibly hypomorphs, or alternatively that the existing *hst* allele and morpholino injections result in hypomorphic or dominant-negative conditions arising from truncated residual protein or lower protein concentration.

The introduction of CRISPR-Cas9 for genome editing has provided the zebrafish field with an easily accessible tool for generating mutant alleles for any gene of choice. Targeted mutagenesis using CRISPR-Cas9 requires careful assessment of targeted candidate gene loci to generate loss-of-function alleles. In contrast to classic forward genetic screens that by definition start from a mutant phenotype linked to a molecular lesion^20^, non-homologous end joining (NHEJ)-based mutagenesis of a candidate locus can result in non-phenotypic lesions. Potential causes of the lack of phenotypes in *de novo* generated mutants include i) translation from downstream start codons, leading to truncated protein products with retained functions that are difficult to assess beforehand; ii) the unpredictable efficiency of nonsense-mediated mRNA decay (NMD) activated in case of premature stop codons; iii) use of alternative, cryptic splice sites to generate functional, translatable mRNA; iv) gene compensation caused by activation of alternative pathways mitigating the phenotype severity. Compensatory mechanisms in mutants have recently been reported in zebrafish for the *egfl7* gene^21^, and the role of compensation in mutant phenotype expressivity and variability in a broader context remains to be assessed.

Of note, the classic *hst* mutant still features detectable *tbx5a* mRNA^7^, and we also detect *tbx5a* transcript by mRNA ISH in *tbx5a^Δ5^-* and *tbx5a^Δ10^-mutant* embryos (Fig. 1X,Y, and data not shown). The *tbx5a^Δ5^*and *tbx5a^Δ10^*lesions are situated in close proximity to the *tbx5a* translation initiation codon; while several possibly initiating ATGs are situated downstream and before the T-box, the amino acid sequence at the N-terminus where our alleles are introduced show conservation from teleosts to mammals (E.C., C.M., data not shown). In addition, frameshift mutations in similar positions within human *TBX5* have been recovered from HOS patients^22^. The full penetrance of concomitant pectoral fin loss and cardiac defects further suggest that no efficient alternative starting codon downstream of the two mutations provides a fully compensating protein product, nor that *tbx5b* would functionally compensate for the function of *tbx5a.* We do acknowledge the possibility that *tbx5b* could act redundant or could compensate for the heartstrings phenotype, clarification of which will require double mutants for both Tbx5-encoding genes in zebrafish.

## CONCLUSIONS

We have generated two new frameshift alleles for *tbx5a* that recapitulate key phenotypes of the published *hst* allele and of morpholino knockdown, with exception of the heartstrings phenotype. While the frameshifts are predicted to form only short out-of-frame proteins, the alleles cannot be conclusively verified as true null alleles. Altogether, our observation underlines the value of analyzing several individual alleles of a candidate gene to assess gene function.

## LIMITATIONS

Due to the unavailability of a Tbx5a-specific antibody or a genetic deletion of the entire tbx5a locus, we cannot verify the absence of Tbx5a protein in our mutants or if the *tbx5a^Δ5^* and *tbx5a^Δ10^* lesions are *bona fide* null alleles. Moreover, we did not assess the possible redundant function of the *tbx5a* paralog *tbx5b,* which is suggested to have a function in pectoral fin specification and heart development^8,9,19^.

## ALTERNATIVE EXPLANATIONS

The absence of the classical *hst* phenotype in our frameshift allele combinations could result from N-terminally truncated protein products from the *tbx5a^Δ5^* and *tbx5a^Δ10^* alleles, which hence would represent hypomorphic alleles. Such products could arise from downstream translation starting off suitable Methionine start codons after the introduced deletion. Nonetheless, the amino acid sequence between the deletions in *tbx5a^Δ5^* and *tbx5a^Δ10^* and the following Methionine codons is conserved across vertebrate species, hinting at functional domains.

## CONJECTURES

Generation of *tbx5b* mutants in the *tbx5a^Δ5^* and *tbx5a^Δ10^* background to discriminate the possible contribution of *tbx5b* to the *tbx5a* null mutant phenotype. Further, complementation analysis with *tbx5a* alleles that feature bigger deletions are required to evaluate if *tbx5a^Δ5^* and *tbx5a^Δ10^* are null alleles or hypomorphs.

## METHODS

Zebrafish*(Daniorerio)* were maintained, collected and staged essentially as described^23,24^. Embryos were raised in temperature-controlled incubators without light cycle at 28°C or as specified in the text. *MO-tbx5a* injection was performed as described^19^ using the translation blocking morpholino sequence *5’-CCTGTACGATGTCTACCGTGAGGC-3’* (Gene Tools). *tbx5a* sgRNA design, synthesis, and injection was performed as described^11^, using the therein published sgRNA sequence sgRNA[tbx5^ccA^] *5’-GGGAGAGTTTTGGAGCC-3’.*

F1 animals were genotyped and alleles recovered as described^11,17^ by amplifying the locus using PCR primers forward *5’-CCACCTGAATAATGTTTGTGC-3’* and reverse *5’-CGGGGATTTGCTGACGGCTG-3’* and sequencing with primer *5’-GCAATCTGAACTAAACTGCA-3’.* The F1 founders harboring the desired alleles were further crossed both into *wildtype* or *RGB* backgrounds. The F2 generation was genotyped again to establish the two strains *tbx5a^Δ5^* and *tbx5a^Δ10^* in *wildtype* and transgenic backgrounds.

The panel plot describing the *tbx5a* alleles was generated using the CrispRVariantsLite software^17^ (http://imlspenticton.uzh.ch:3838/CrispRVariantsLite/) from Sanger sequencing data of the alleles.

Each clutch was assessed for mortality at 24hpf and phenotype percentages at 3dpf were calculated on living embryos. Statistical parameters and analysis were assayed with GraphPad Prism 6, Student’s t-test was performed on at least three independent clutches for each allele combination and a p-value less than 0.0001 was considered significant. Embryos for imaging were raised in E3 with 1-phenyl 2-thiourea (PTU) 200μM final and treated at the moment of imaging with tricaine pH=7 3% final and 20mM 2,3-butanedione monoxime (BDM, Sigma Aldrich) to temporarily stop heartbeat.

The triple-transgenic reporter transgenic strain RGB (Red, Green, Blue) was generated by crossing together the transgenic insertions *Imo2:loxP-dsRED2-loxP-EGFP^u^, drl:EGFP*^15^, and *myl7:loxP-AmCyan-loxP-ZsYellow*^16^ and propagation by repeated in- and out-crosses followed by thorough screening.

Fluorescent stereomicroscopy of *RGB* of *wildtype, tbx5a^Δ5^,tbx5a^Δ10^* and *tbx5a^Δ5/Δ10^* was performed with a Leica M205 FA equipped with 1x magnification lens at 100x magnification. Samples for SPIM imaging were mounted in 0.5% low melting agarose and imaged with a 20x magnification lens plus a 10x illumination source in a Zeiss Z.1 (ZMB, UZH). Image stacks were processed with ImageJ1.50d and Gimp2.8.

The *in situ* probes for *versican a* (vcana)^7,10^ and *tbx5a* were generated cloning-free by adding the *T7* promoter sequence to the reverse primer; primers used were *(T7* promoter underlined):

- *vcana fw 5’-AGCTGACCCAGACATTGAGG-3’,*
- *vcana rev 5<u>-TAATACGACTCACTATAGGG</u>TCTTGCAGGTGAAAGTGAGG-3’,*
- *tbx5a fw 5-GGCGGACAGTGAAGACACCTT-3’,*
- *tbx5a rev 5<u>-TAATACGACTCACTATAGGG</u>CATGATGCTGACGCTGTGCA-3’.*

The PCR templates were purified with a QIAquick Gel Extraction kit. The *in situ* probes were transcribed using T7 RNA Polymerase (Roche) and DIG RNA labeling Mix (Roche) according to manufacturer instructions. Probe quality was assessed via MOPS gel electrophoresis under denaturizing conditions. Whole-mount *in situ* hybridization (ISH) was performed using standard protocols with bleaching post-hybridization^19,25^.

## ACKNOWLEDGMENTS

The authors like to thank Dr. Nadia Mercader for insightful discussions, Dr. Stephan Neuhauss and Kara Dannenhauer for assistance with zebrafish husbandry, the ZMB team at UZH for critical imaging support, and all members of the Mosimann lab for continued support and input.

### FUNDING STATEMENT

This work was supported by the Canton of Zürich and a project grant from the Swiss Heart Foundation; a Swiss National Science Foundation (SNSF) professorship (PP00P3_139093) and a Marie Curie Career Integration Grant from the European Commission (CIG PCIG14-GA-2013-631984) to C.M.; a UZH URPP "Translational Cancer Research” Seed Grant to A.B.; and a SNSF R’Equip Grant (316030_150838/1).

### ETHICS STATEMENT

Zebrafish for embryo production were kept in the UZH Irchel campus zebrafish facility (Tierhaltungsnummer 150). All experiments use zebrafish embryos up to 5 dpf/120 hours. Experiments with zebrafish embryos up to the age of 120 hours are not considered animal experiments by Swiss law (Art. 112 Bst. d) and the revised Directive 2010/63/EU (outlined in Strähle et al.,Reprod Toxicol. 2012 Apr;33(2):128-32.), as confirmed by the animal ethics office at UZH.

